# SC-GAN: 3D self-attention conditional GAN with spectral normalization for multi-modal neuroimaging synthesis

**DOI:** 10.1101/2020.06.09.143297

**Authors:** Haoyu Lan, the Alzheimer Disease Neuroimaging Initiative, Arthur W Toga, Farshid Sepehrband

## Abstract

Image synthesis is one of the key applications of deep learning in neuroimaging, which enables shortening of the scan time and/or improve image quality; therefore, reducing the imaging cost and improving patient experience. Given the multi-modal and large-scale nature of neuroimaging data, the synthesis task is computationally challenging. 2D image synthesis networks do not take advantage of multi-dimensional spatial information and the 3D implementation has dimensionality problem, negatively affecting the network reliability. These limitations hinder the research and clinical applicability of deep learning-based neuroimaging synthesis. In this paper, we proposed a new network that is designed and optimized for the application of multi-modal 3D synthesis of neuroimaging data. The network is based on 3D conditional generative adversarial network (GAN), and employs spectral normalization and feature matching to stabilize the training process and ensure optimization convergence. We also added a self-attention module to model relationships between widely separated voxels. The performance of the network was evaluated by predicting positron emission tomography (PET) images, Fractional anisotropy (FA) and mean diffusivity (MD) maps from multi-modal magnetic resonance images (MRI) of 265 and 497 individuals correspondingly. The proposed network, called self-attention conditional GAN (SC-GAN), significantly outperformed conventional 2D conditional GAN and the 3D implementation, enabling robust 3D deep learning-based neuroimaging synthesis.

## Introduction

Medical image synthesis is a technique to generate new parametric images from other medical image modalities that contain a degree of similarity or mutual information. Medical image synthesis can be used for a number of valuable applications, including shortening imaging time, data augmentation, enabling low dose contrast administration and even image enhancement (Hiasa et al., 2018; Nie et al., 2017; Roy et al., 2014; Shin et al., 2018; Wang et al., 2018; Yi et al., 2019). For the last five years, the power of image synthesis has been proven in computer vision and image processing fields. In particular, generative adversarial networks (GAN) (Goodfellow et al., 2014) has been shown to be an effective and reliable technique for image synthesis (Huang et al., 2018). Variants of GAN like Conditional GAN (Mirza and Osindero, 2014) and Cycle GAN (Zhu et al., 2017) also have been proposed to generalize GAN to different tasks and circumstances, including medical image synthesis.

Medical image synthesis with deep convolutional neuronal networks are often implemented using encoder-decoder networks, GAN or its variants. For example, Nie *et al* (Nie et al., 2018) proposed a deep convolutional adversarial network to synthesize Computer Tomography (CT) images from magnetic resonance images (MRI), and also to synthesize 7T MRI from 3T images. Chen *et al* (Chen et al., 2019) implemented encoder-decoder convolutional neural network to synthesis positron emission tomography (PET) from ultra-low dose PET and MRI. Ouyang *et al* (Ouyang et al., 2019) used conditional GAN with task specific perceptual loss to synthesize PET from ultra-low dose PET. These techniques used 2-dimensional (2D) or 2.5D network implementations. For a 2D network implementation, image slices along one of 3D anatomical planes (i.e. axial, coronal and sagittal) are trained independently and then combined or ensembled in decoding step. Employing a 2D approach on a 3D data is suboptimal and inefficient because it does not incorporate the 3D spatial information in the image, and/or it requires multiple independent implementation of a same network along different image axes.

3D networks were proposed to address the limitations of the 2D and 2.5D networks for the purpose of image synthesis. Wang *et al* (Wang et al., 2019) proposed a 3D conditional GAN network for PET synthesis from low dose input, which resulted to improved network performance in comparison with the 2D implementation. Liu *et al* (Liu, 2019) showed that 3D GAN performance can be improved further by incorporating an attention gate module to generate synthesis results, which they used as the input of a segmentation task. Given that the aim of image synthesis is to generate a new image from the existing images of the same individual, we anticipate that a self-attention module could further improve the performance of GAN. The 3D implementation of self-attention GAN however, with no specific modification/addition to network elements and optimizers, creates inconsistency problem due to the large differences of feature distributions (Wang et al., 2019), negatively affecting the network reliability and sometimes fails to converge. In order to improve GAN performance and to address these limitations, we developed a new 3D GAN and optimized it for neuroimaging synthesis.

Proposed 3D Self-attention Conditional GAN (SC-GAN) is constructed as follow: First, we extended 2D conditional GAN into 3D conditional GAN. Then, we added 3D self-attention module to 3D conditional GAN to generate 3D images with preserved brain structure and reduced blurriness within the synthesized images. We also introduced spectral normalization (Miyato et al., 2018), feature matching loss (Wang et al., 2018) and brain area root mean square error (RMSE) loss to stabilize training and prevent overfitting. SC-GAN is an end-to-end medical image synthesis network that can be applied on high-resolution input images (e.g. 256 x 256 x 256). SC-GAN can also be applied on multi-modal input data and is designed using 3D convolutional layers.

The novelties and contributions of this technique are as follows.

I. For the first time, combining 3D self-attention module into 3D conditional GAN to generate high accuracy synthesis results with stable training process. A smooth training was achieved by using a series of stabilization techniques and modified loss function.
II. SC-GAN was tested on multiple datasets across different synthesis tasks and enables multi-model input, which can be generalized to other image synthesis applications.
III. SC-GAN source code is made available at https://github.com/Haoyulance/SC-GAN

### Theory and Method

Here we introduce the 3D Self-attention Conditional GAN (SC-GAN) theory and the mathematical formulation of its components.

### 3D conditional GAN

For the main body of the SC-GAN, we used conditional GAN, which is shown to be the optimum choice of GAN for medical image synthesis and reconstruction with paired images (Wang et al., 2019)(Ouyang et al., 2019)(Li et al., 2020)(Zhao et al., 2020). SC-GAN was then designed by adding additional modules to conditional GAN that are described in detail in the next sections. This section describes the conditional GAN, which was also used as the baseline for evaluating the SC-GAN.

In an unconditional GAN (Goodfellow et al., 2014), the generator learns the mapping from the latent space to target image space by adversarial learning to generate the fake outcome, without any label specify. Conditional GAN on the other hand learns to generate the outcome using a specific condition, allowing the application of supervised learning for image-to-image generation. Therefore, when ground truth data is available for training, conditional GAN is a powerful network to do image translation. Conditional GAN uses below loss function:

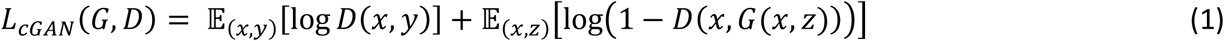

where *x* is the input image and also the condition image, *y* is the ground truth image and *z* is the Gaussian noise. Notice *z* is the sample in the latent space for unconditional GAN to generate stochastic results. As for image to image translation, condition image *x* has enough variance so that generator would easily learn to ignore *z* (Isola et al., 2017), (Ouyang et al., 2019). Therefore, in conditional GAN noise *z* is no longer provided to the generator and the loss function is formulated as:

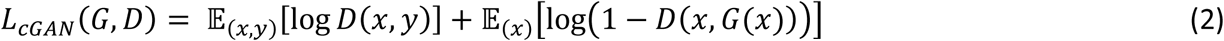

We adopted pix2pix (Isola et al., 2017), a variant network of 2D conditional GAN, as the network structure of 3D conditional GAN. in our experiment, 3D conditional GAN has 8 layers generator, similar to U-net (Ronneberger et al., 2015), and uses PatchGAN classifier(Isola et al., 2017) as the discriminator. The objective function is:

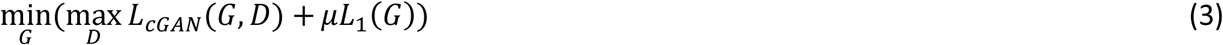

Where *L*_1_(*G*) = 𝔼_(*x,y*)_(‖*y* − *G*(*x*)‖_1_) is the *L*_1_ loss between the ground truth and generated image and *μ* is the regularization term for the *L*_1_ loss.

Generator’s optimization aims to minimize the objective function. Only generator’s weights are updated in each iteration of the optimization. Discriminator’s optimization aims to maximize the objective function and therefore only discriminator’s weights are updated in each iteration. Generator and discriminator forward and backward propagate alternately till the training process reaches Nash equilibrium and network converges (Nash, 1950).

### Feature matching loss

To stabilize the training, we incorporated a feature matching loss (Wang et al., 2018). Feature matching loss is described as follow:

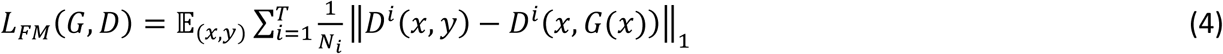

where *D*^*i*^ is the *i*_*th*_ layer’s feature map; *T* is the total number of layers of discriminator and *N*_*i*_ is the number of elements in *i*_*th*_ layer’s feature map.

Feature matching loss was added only to the generator loss, because only the *L*_*FM*_ is required to be minimized at generator’s optimization. The objective function with feature matching loss is:

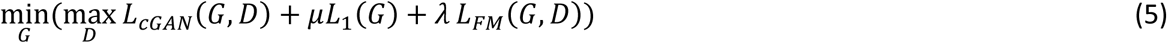

where regularization term (*λ*) controls the importance of the feature matching loss.

### Brain area RMSE loss

Error calculation was done on brain voxels and the background was excluded. We calculated root mean square error (RMSE) between masked *G* and masked *y*, then added the RMSE to the generator loss. We obtained the brain area (*mask*_*y*_) from the ground truth *y*, then, which was used to calculate brain area RMSE (B-rmse) loss:

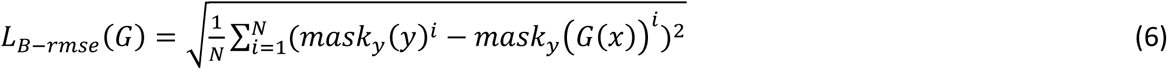

where *mask*_*y*_(*y*)^*i*^ is the *i*_*th*_ voxel of *mask*_*y*_(*y*) and *N* is the number of total voxels. Objective function of B-rmse loss is:

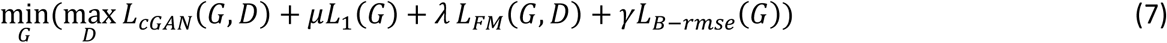

where *γ* controls the regularization term for the brain area rmse loss.

In the ablation study, we found that B-rmse loss contributed to the improvement of the network performance and improved the synthesis accuracy. Notice that B-rmse loss is not the only loss for the generator, there are combination of *L*_1_ loss, B-rmse loss and feature matching loss for generator. *L*_1_ loss focuses on the difference of whole output and target and B-rmse loss focuses on the only brain area’s difference of output and target.

### 3D self-attention

Self-attention allows GAN to efficiently model relationships between widely separated spatial regions (Zhang et al., 2018), so that generated images contain realistic details. The image feature map *x ϵ R*^*C*×*h*\**w*\**d*^ from one intermediate hidden layer of 3D cGAN was transformed into 2 feature spaces *f*(*x*) = *W*_*f*_*x* and *g*(*x*) = *W*_*g*_*x* to calculate the attention. Then, the third feature space *h*(*x*) = *W*_*h*_*x* was used to calculate attention feature map. Since the purpose of utilizing self-attention is to measure the similarity of each voxel with target voxel, we used the similarity scores (attentions) as weights to calculate the weighted sum represent of each target voxel. 3D self-attention module structure is presented in **Figure 1**.

**Figure 1.**
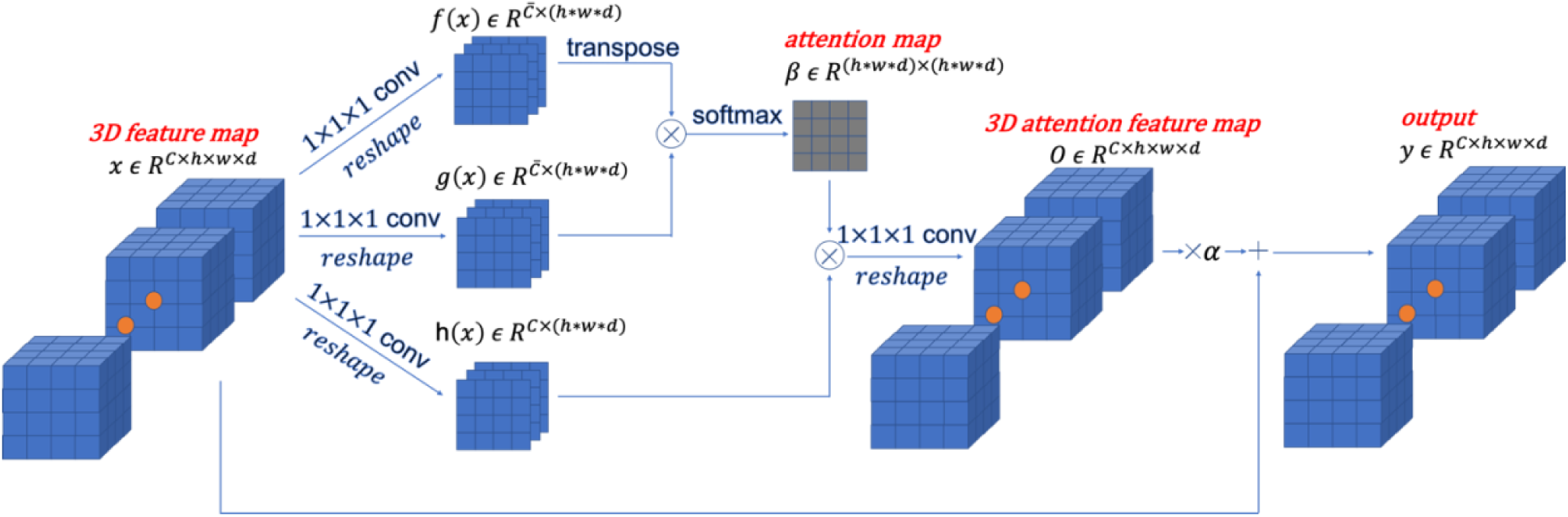
3D self-attention module. This Figure is a schematic view of the self-attention module of SC-GAN. The first layer represents the input data. Attention map exploit the similarity of each pair of convolved images and combine it with the input data to generate the output of the self-attention module.

Similarity score (attention) was calculated as follow:

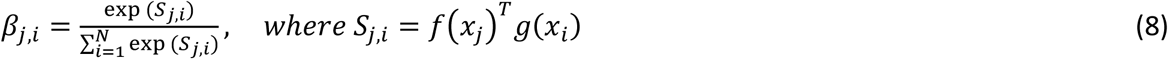

in which, *β*_*j,i*_ is voxel *j*’s attention to voxel *i*. We then calculated attention feature for each voxel *j* by:

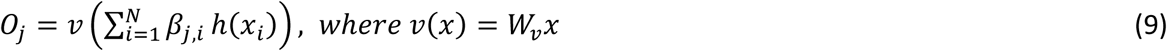

The final output of attention layer is:

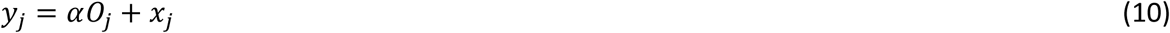

In the above formulations

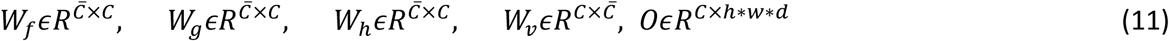

*W*_*f*_, *W*_*g*_, *W*_*h*_, *W*_*v*_ are learned weight matrices by 1 × 1 × 1 3D convolutions; *C* is the number of original channels; 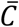 equals to *C*/8 for memory efficiency; *h* * *w* * *d* is the number of voxels in one feature map; *α* is a learnable scalar and it is initialized as 0.

In our network, self-attention is implemented in both generator and discriminator as shown in **Figure 2**. Generator for conditional GAN is the same as U-net (Ronneberger et al., 2015) structure. When comparing our results with U-net, we added self-attention at both encoder and decoder of U-net to improve the synthesis performance.

**Figure 2.**
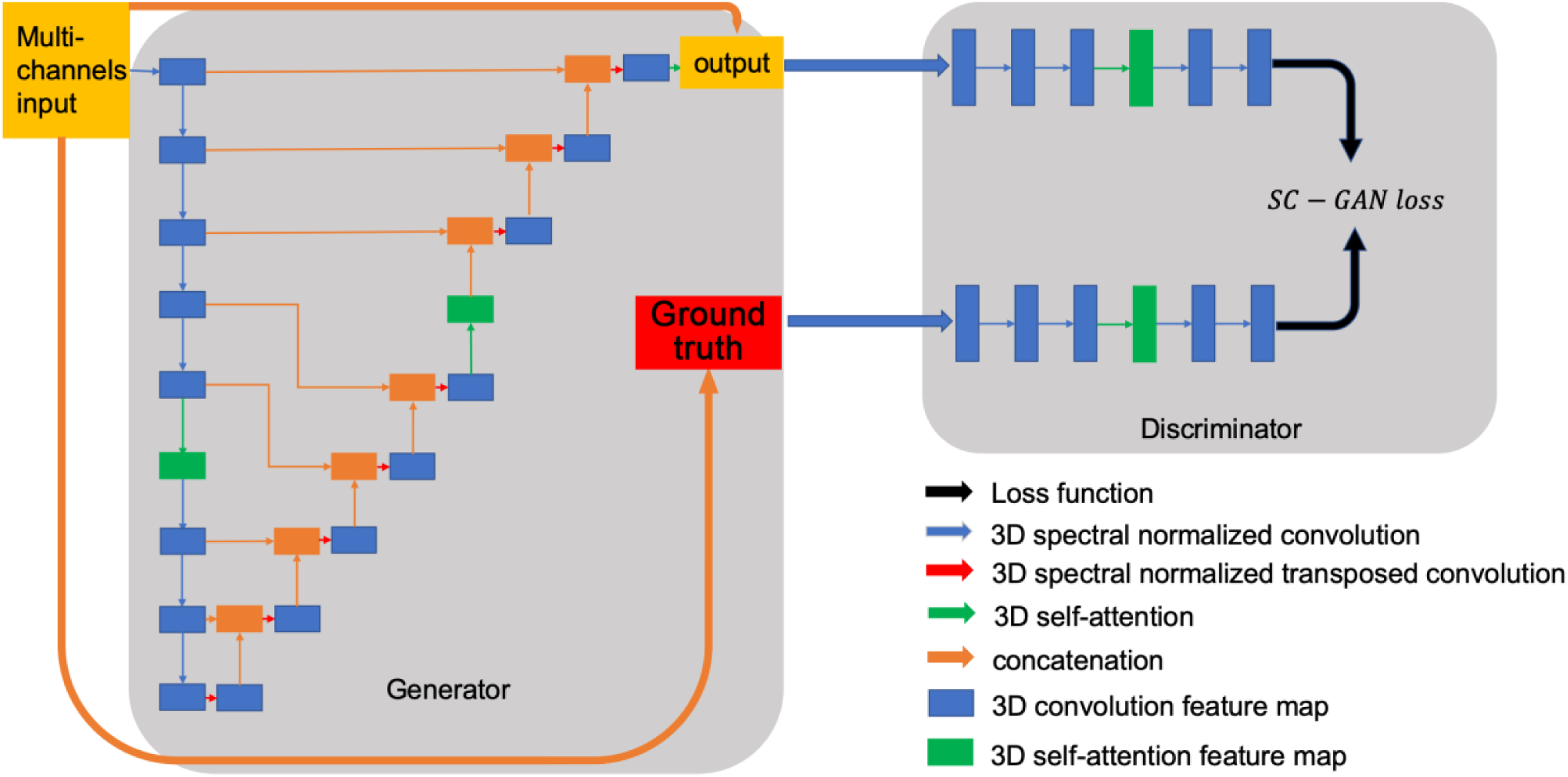
SC-GAN structure with 3D self-attention module. The network structure of SC-GAN constitutes of two parts generator and discriminator. The generator is a Unet like 8 layers encoder-decoder with 3D self-attention module in the middle of encoder and decoder. The discriminator is a 5 layers patch GAN with 3D self-attention. Self-attention module empowers the both generator and discriminator in the adversarial learning strategy.

### Spectral normalization

Spectral normalization is implemented in each layer *g*: *h*_*in*_ → *h*_*out*_ of the neural networks to normalize the weight matrix between two connected layers by controlling the Lipschitz constant. By definition, Lipschitz norm ‖*g*‖_*Lip*_ = *sup*_*h*_*σ*(∇*g*(*h*)), where *σ*(·) is the spectral norm (the largest singular value).

Suppose a neural network *f*(*x, W, a*) = *W*^*L*+1^*a*_*L*_(*W*_*L*_(*a*_*L*−1_(*W*^*L*−1^(… *a*_1_(*W*^1^*x*) …)))), where {*W*^1^,*W*^2^, …, *W*^*L*+1^} is the weights set, {*a*_1_,*a*_2_, …, *a*_*L*_} is the element-wise non-linear activation functions. For the linear layer *g*(*h*) = *Wh*, the norm is given by:

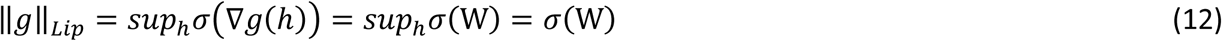

If the Lipschitz norm of the activation function ‖*a*_*L*_‖_*Lip*_ is equal to 1, based on the Cauchy-Schwarz inequality ‖*g*1 ∘ *g*2‖_*Lip*_ ≤ ‖*g*1‖_*Lip*_ · ‖*g*2‖_*Lip*_, the following bound can be derived:

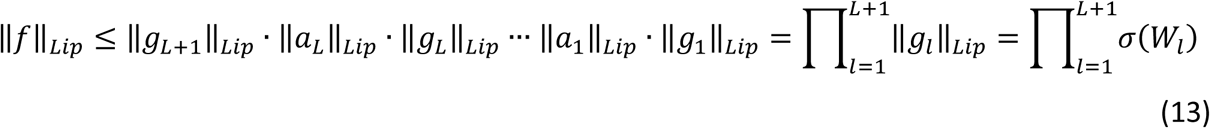

The spectral normalization normalizes the spectral norm of the weight matrix *W*_*l*_ to get *W*_*SN*_ = *W*_*l*_/*σ*(*W*_*l*_). Thus, if *W*_*l*_ is normalized as *W*_*SN*_, then 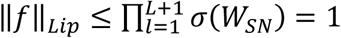 which means ‖ *f*‖_*Lip*_ is bounded by 1. Miyato *et al* (Miyato et al., 2018) have shown the importance of Lipschitz continuity assuring the boundness of statistics. We utilized Spectral normalization in both generator and discriminator of SC-GAN.

### Regularization

In order to prevent overfitting, we added L2 norm regularizations to generator and discriminator, resulting to a final objective function of:

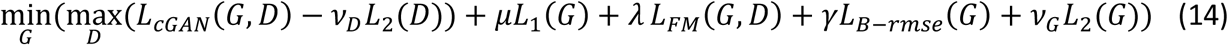

where *v*_*D*_ and *v*_*G*_ control the importance of *L*_2_ regularization. Since during the training process we minimize the negative discriminator loss for the discriminator training, the above objective function uses −*v*_*D*_*L*_2_(*D*) to regularize discriminator. Note that *L*_2_(*D*) and *L*_2_(*G*) are the constraints on trainable values of discriminator and generator, however, *L*_1_(*G*) is the *L*_1_ distance between generated output and ground truth.

## Experiments

### Study data

Data used in the preparation of this article were obtained from the Alzheimer’s Disease Neuroimaging Initiative 3 (ADNI-3) database (http://adni.loni.usc.edu) (Weiner et al., 2017). We downloaded MRI and PET data from ADNI-3 participants. All available from ADNI-3 at the time of this study were used for this study (ADNI-3 is an ongoing project). For PET synthesis task 265 images were used (training set = 207, testing set = 58). For FA and MD synthesis tasks 497 images were used (training set = 398, testing set = 99). For MRI, T1-weighted (T1w) and fluid-attenuated inversion recovery (FLAIR) structural magnetic resonance image (MRI) and diffusion-weighted MRI were used. For PET, we used amyloid PET data. For PET synthesis, dataset with complete T1w, FLAIR and amyloid PET sessions with acceptable quality, based on ADNI guidelines were included in the analysis. For diffusion-weighted MRI synthesis, dataset with complete T1w, FLAIR and diffusion-weighted MRI sessions were used (all images were visually inspected).

### MRI data collection and preprocessing

MRI imaging of the ADNI-3 was done exclusively on 3T scanners (Siemens, Philips, and GE) using a standardized protocol. 3D T1w with 1 mm^3^ resolution was acquired using an MPRAGE sequence (on Siemens and Philips scanners) and FSPGR (on GE scanners). For FLAIR images, a 3D sequence with similar resolution as T1w images was acquired, which provided the opportunity for accurate intrasubject intermodal co-registration. MPRAGE T1w MRI scans were acquired using the following parameters: TR = 2300 ms, TE = 2.98 ms, FOV = 240 × 256 mm^2^, matrix = 240 × 256 (variable slice number), TI = 900 ms, flip angle = 9, effective voxel resolution = 1 × 1 × 1 mm^3^. The FSPGR sequence was acquired using sagittal slices, TR = 7.3 ms, TE = 3.01 ms, FOV = 256 × 256 mm^2^, matrix = 256 × 256 (variable slice number), TI = 400 ms, flip angle = 11, effective voxel resolution = 1 × 1 × 1 mm^3^. 3D FLAIR images were acquired using sagittal slices, TR = 4,800 ms, TE = 441 ms, FOV = 256 × 256 mm^2^, matrix = 256 × 256 (variable slice number), TI = 1650 ms, flip angle = 120, effective voxel resolution = 1 × 1 × 1.2 mm^3^.

T1w preprocessing and parcellation was done using the FreeSurfer (v5.3.0) software package, which is freely available (Fischl, 2012), and data processing using the Laboratory of Neuro Imaging (LONI) pipeline system (http://pipeline.loni.usc.edu) (Dinov et al., 2010, 2009; Moon et al., 2015; Torri et al., 2012), similar to (Sepehrband et al., 2018; Sta Cruz et al., 2019). Field corrected, intensity normalized images were filtered using non-local mean filtering to reduce the noise, and the outputs were used for the analysis. FLAIR images of each individuals were corrected for non-uniform field inhomogeneity using N4ITK module (Tustison et al., 2010) of Advanced Normalization Tools (ANTs) (Avants et al., 2009). FLAIR images were then co-registered to T1w images using *antsIntermodalityIntrasubject* ANTs module.

Diffusion MRI is a quantitative modality and contain microstructural information about brain tissue (Le Bihan et al., 2001; Sepehrband et al., 2017, 2015). Therefore, it was used as a challenging synthesis target from T1 and FLAIR, which are mainly qualitative maps. Diffusion MRI data was acquired using the following parameters: 2D echo-planar axial imaging, with sliced thickness of 2mm, in-plane resolution of 2mm^2^ (matrix size of 1044 x 1044), flip angle of 90°, 48 diffusion-weighted images with 48 uniformly distributed diffusion-encodings with b-value=1000 s/mm^2^ and 7 non-diffusion-weighted images. Diffusion MRI preprocessing and diffusion tensor imaging (DTI) fitting were performed were as described in (Sepehrband et al., 2019b, 2019a). In brief, images were corrected for eddy current distortion and for involuntary movement, using FSL TOPUP and EDDY tools (Andersson et al., 2012, 2003). DTI was then fitted to diffusion data using Quantitative Imaging Toolkit (Cabeen et al., 2018). Fractional anisotropy (FA) and mean diffusivity (MD) maps were used for the synthesis task.

### PET data collection and preprocessing

Amyloid PET analysis was performed according to UC Berkeley PET methodology for quantitative measurement (Baker et al., 2017; Landau et al., 2015, 2014; Schöll et al., 2016). Participants were imaged by Florbetapir (^18^ F-AV-45, Avid), or ^18^ F-Florbetaben (NeuraCeq, Piramal). Six five-minute frames of PET images were acquired 30 to 60 minutes post injection. Each extracted frame is co-registered to the first extracted frame and then combined into one image, which lessens the subject motion artifacts. The combined image had the same image resolution of the original PET image (2mm isotropic voxels). All PET images were co-registered on T1w MRI. Quantitative measurement was done based on Standard Uptake Value ratio (SUVR). The brain mask was, obtained from T1w analysis was applied on co-registered. T1w, FLAIR and PET images. Examples of a set of input and target images are presented in **Figure 3**.

**Figure 3.**
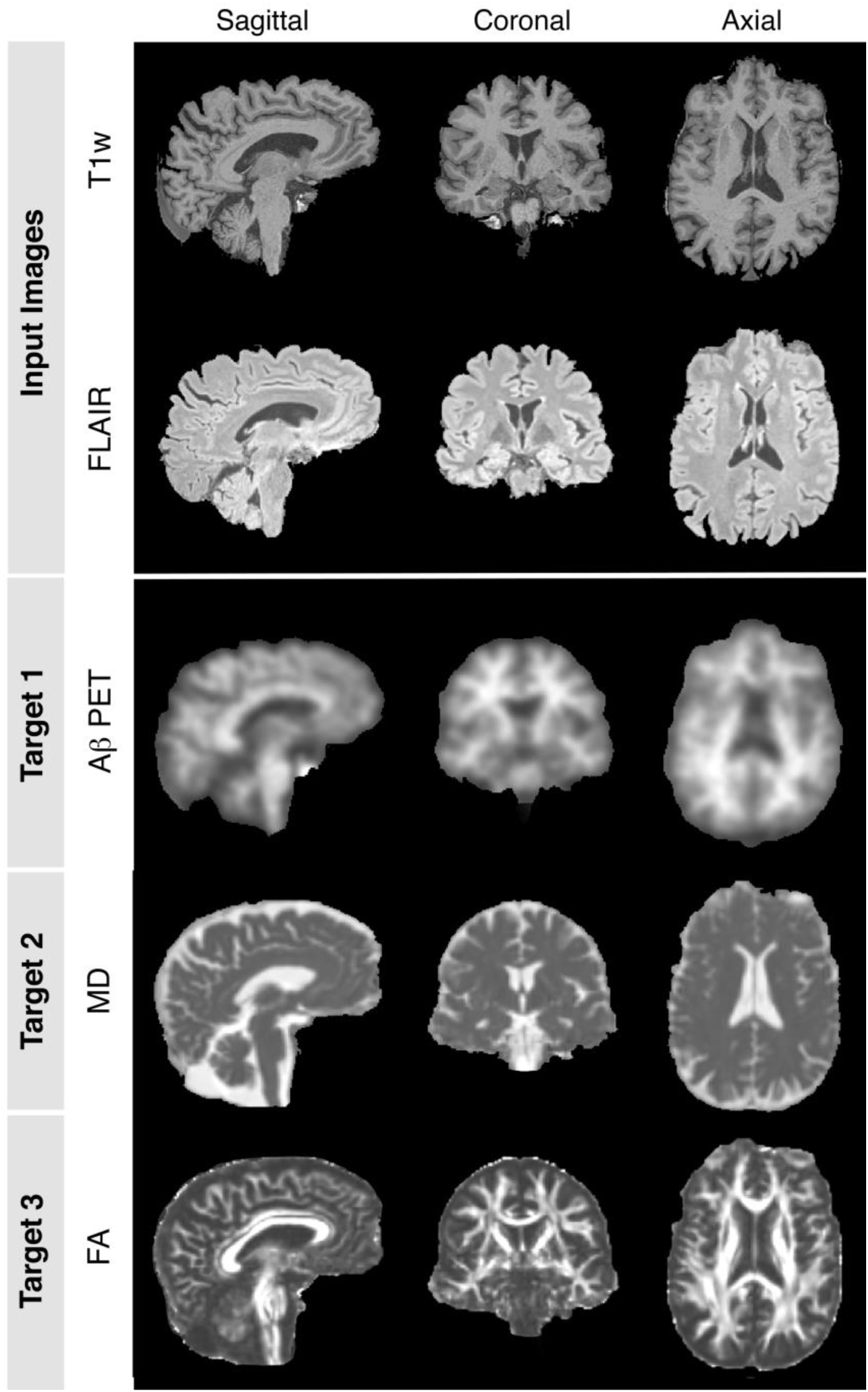
Multi-modal (multi-channel) input. Examples of different neuroimaging data from single individual are presented. T1-weighted (T1w) and fluid-attenuated inversion recovery (FLAIR) were used as input for different synthesis tasks. For each the study tasks a different target was used, which are shown as outputs 1-3: Mean Diffusivity (MD), Fractional Anisotropy (FA) and Amyloid-beta Positron Emission Tomography (Aβ-PET). Data were preprocessed and co-registered (see method section for detail), are shown from three anatomical views (from left to right: axial, coronal and sagittal).

### Implementation, baseline models

In order to rigorously assess the performance of the SC-GAN, we have compared it with current well-developed medical image synthesis networks, including: 2D cGAN, 3D cGAN and Attention cGAN (Att cGAN). 2D cGAN was adopted from Ouyang *et al* (Ouyang et al., 2019), which proposed it for PET synthesis task. 3D cGAN was firstly proposed by Wang *et al* (Wang et al., 2019) for PET image synthesis from low dose PET images. Attention cGAN was designed based on the attention module proposed by Oktay *et al* (Oktay et al., 2018), which incorporated the 3D attention module in the U-net architecture for the application of pancreas segmentation (assisted by image synthesis task). The same 3D attention module was also adopted by Liu *et al*(Liu, 2019) in Cycle-GAN medical image synthesis network. For a fair comparison, we incorporated aforementioned 3D attention module in conditional GAN, here referred to as Att-GAN, and compared it with SC-GAN. Note that the self-attention module has a different mechanism compared with attention module. Unlike the attention module, the self-attention exploits the dependencies of each pair of positions in the same feature map to get attention matrix, then use attention matrix to reconstruct representation and combine it with the same input data. All 3 baseline models and SC-GAN were implemented using TensorFlow (1.12.2) and deployed training on an NVIDIA GPU cluster equipped with eight V100 GPUs (Cisco UCS C480 ML). All four sets of results are used to analyze and compare different networks’ performance.

### Image preparation prior to training

For the PET synthesis task, 207 pairs of T1W and FLAIR images were used as training data and 58 pairs of T1w and FLIAR images were used as test data. For the DTI-MD and DTI-FA synthesis tasks, 398 pairs of T1W and FLAIR images were used as training data and 99 pairs of T1w and FLIAR images were used as test data. PET and DTI were upsampled to have the same resolution as the T1 and FLAIR, i.e. 256 x 256 x 256. We implemented Z-score normalization for all four tasks, then applied min-max rescaling to scale the voxels’ intensity between 0 to 1 prior to the training. Synthesis methods could be improved with intensity normalization, but are robust to the choice of the normalization (Reinhold et al., 2019).

### Training and testing

2D cGAN was implemented similar to (Ouyang et al., 2019). We utilized pix2pix structure (U-net generator and patch GAN discriminator) with feature matching loss and regularization. 3D cGAN was implemented similar to (Wang et al., 2019) and Att cGAN was implemented similar to (Liu, 2019; Oktay et al., 2018). We performed 5-fold cross validation during the hyperparameter tuning phase for all four networks to get the optimum hyperparameters. SC-GAN network architecture is illustrated in **Figure 2** and the loss function formulation was described in equation 14. The optimum result was obtained with the following hyperparameters: *μ* = 200, *γ* = 200, *λ* = 20, *v*_*G*_ = 0.001, *v*_*D*_ = 0.001, batch size=1. Learning rate starts as 0.001 and cosine decay was used to continuously shrink the learning rate during the training process.

### Evaluation criteria

Three image quality metrics were used to evaluate the performance of the synthesis task: normalized root mean square error (NRMSE), peak signal-to-noise ratio (PSNR) and structural similarity (SSIM). NRMSE reflects the normalized error without being affected by the range of the voxel values. Thus, NRMSE could be used to compare the performances of the network on different tasks directly. To enable a direct comparison between 2D cGAN and 3D networks, we evaluated the 3D output of the 2D network directly.

### Ablation study

In order to analyze the contribution of each component of SC-GAN, we performed an ablation study. Five ablation tests were conducted for the proposed network, namely: SC-GAN 1) without self-attention module, 2) without adversarial learning, 3) without brain area rmse loss, 4) without spectral normalization, and 5) without feature matching loss.

### Evaluating synthesized PET

A secondary analysis was performed to compare SC-GAN results against *ground truth* PET. Amyloid-b (Aβ) uptake were estimated from PET and synthesized PET. The Aβ uptake values were then compared across clinically relevant regions. While the focus of the study was on proposing and optimizing a neuroimage synthesis technique, this evaluation was performed to examine whether the PET synthetization from MRI can substitute the PET imaging. Standard uptake value ratio (SUVR) of the Aβ were calculated across subcortical and cortical regions of 10 randomly selected individuals from ADNI-3 cohort. SUVR values of 110 regions per participants were compared between PET and synthesized PET. SUVRs across these regions of interest were derived using the Desikan-Killiany atlas, which were parcellated on T1w images using FreeSurfer pipeline, as explained in *B. MRI data collection and preprocessing* section. PET images that were used for training were normalized using min-max normalization approach. Therefore, test PET images were also normalized using the same approach before comparison.

## Results

The learning curves of the GANs that were used for PET, FA and MD synthesis tasks are presented in the **Figure 4.** Learning curves demonstrate network performance across training epochs. Average performance of applying the trained network on the test data is presented in **Figure 5**, and the qualitative assessments are presented in **Figures 6-8**.

**Figure 4.**
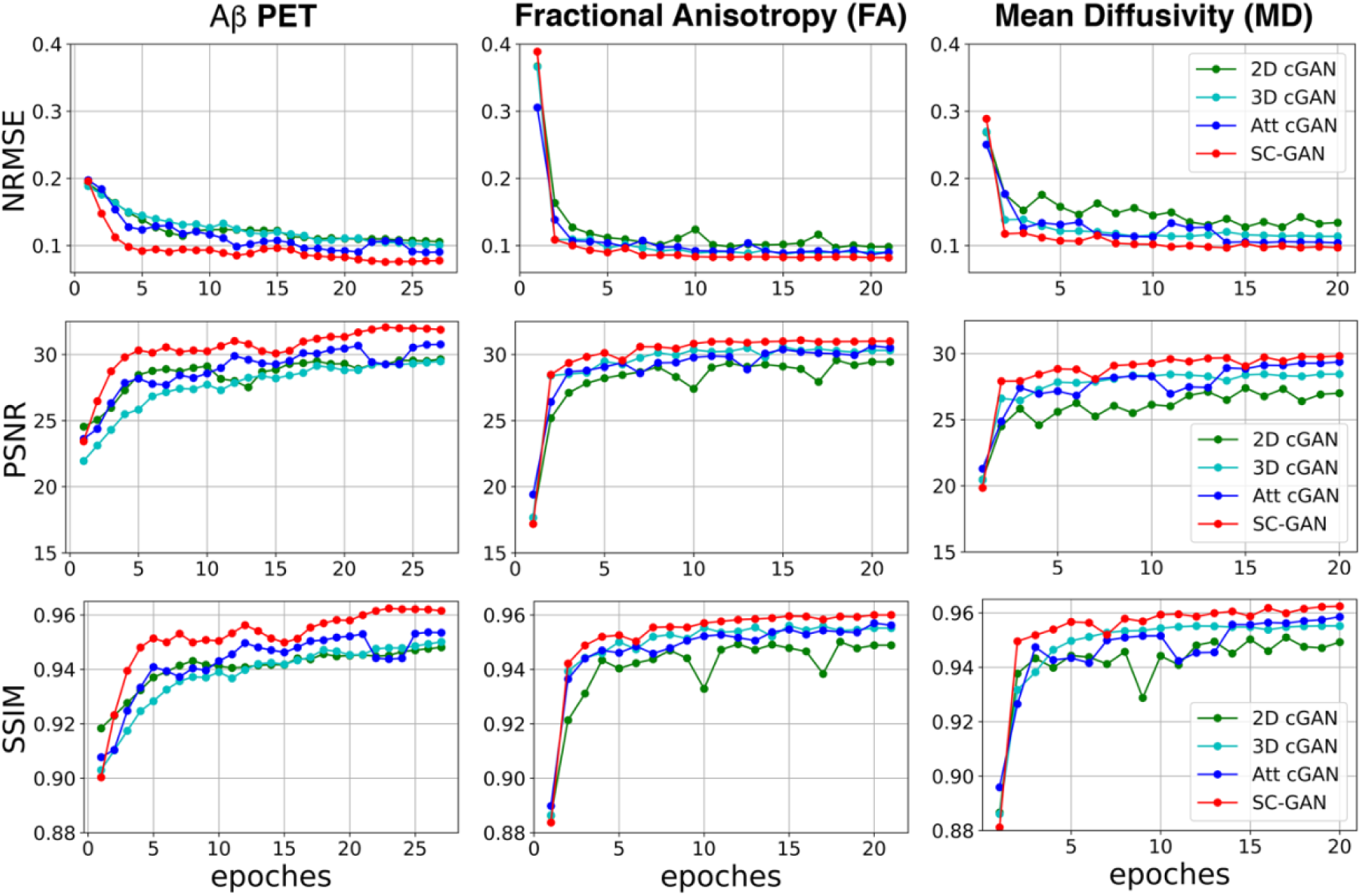
Learning curves SC-GAN compared to other Synthesis GANs across different taks. Plots demonstrate learning curves of four CNN networks that were evaluated in this study: 2D GAN, 3D GAN, 3D cGAN with Attention gate (Att cGAN) and SC-GAN. T1w and FLAIR were used for three tasks: 1) synthesizing Amyloid-beta PET (n=242, **first column**); 2) synthesizing fractional anisotropy (n=480, **second column**); 3) synthesizing mean diffusivity (n=480, **third column**). Three different evaluation metrics were used: **First row** shows normalized root mean square error (NRMSE); **Second row** shows peak signal-to-noise ratio (PSNR); **Third row** shows structural similarity (SSIM). Note that all networks reached their plateau around epoch=20.

**Figure 5.**
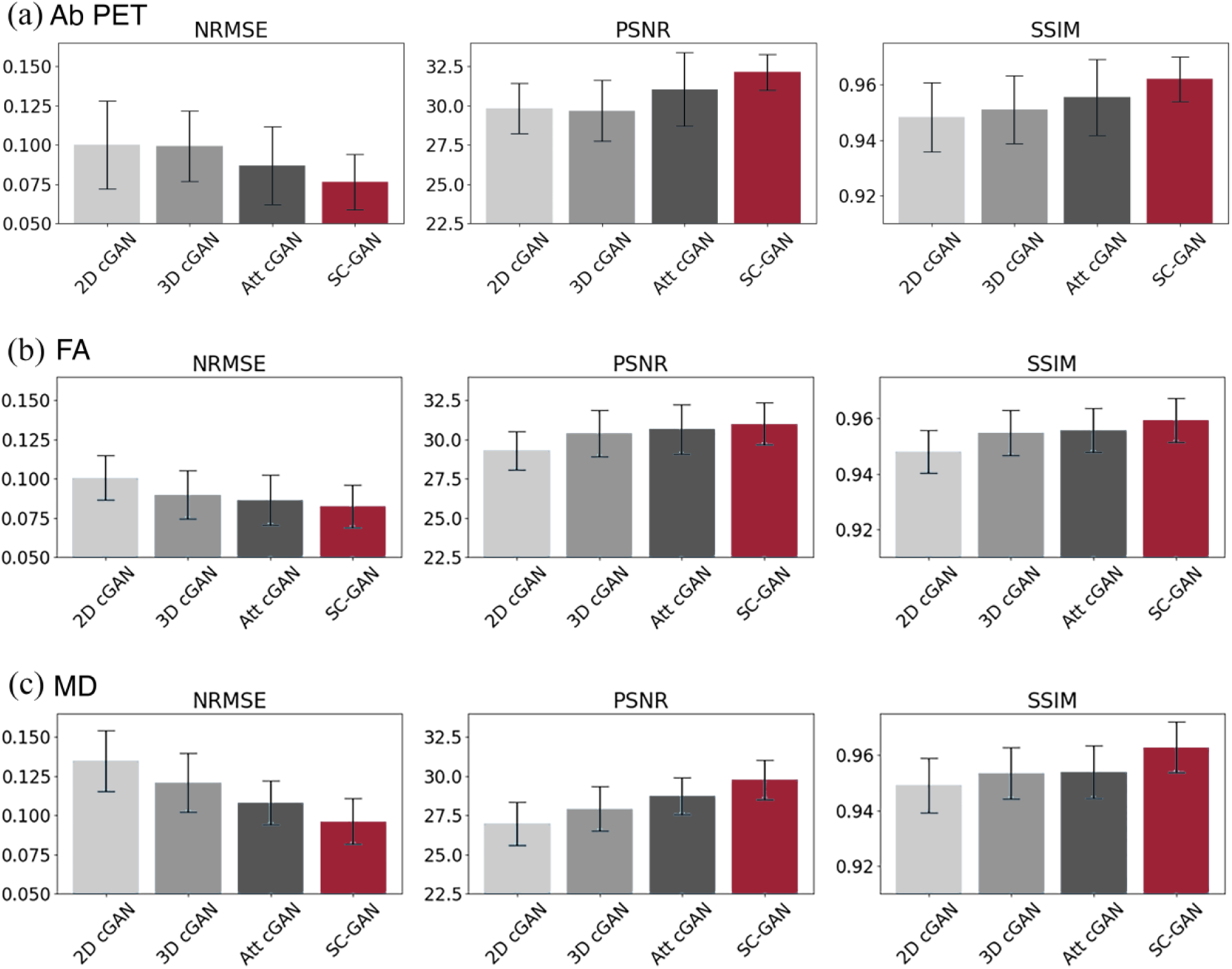
Image quality metrics on test data across different tasks. Bar charts demonstrate normalized mean square error (NRMSE), peak signal-to-noise ratio (PSNR) and structural similarity (SSIM) among test images after the networks reached the plateau and the hyperparameters were optimized. T1w and FLAIR were used for three tasks: 1) synthesizing Amyloid-beta PET (n=242, **A**); 2) synthesizing fractional anisotropy (n=480, **B**); 3) synthesizing mean diffusivity (n=480, **C**).

**Figure 6.**
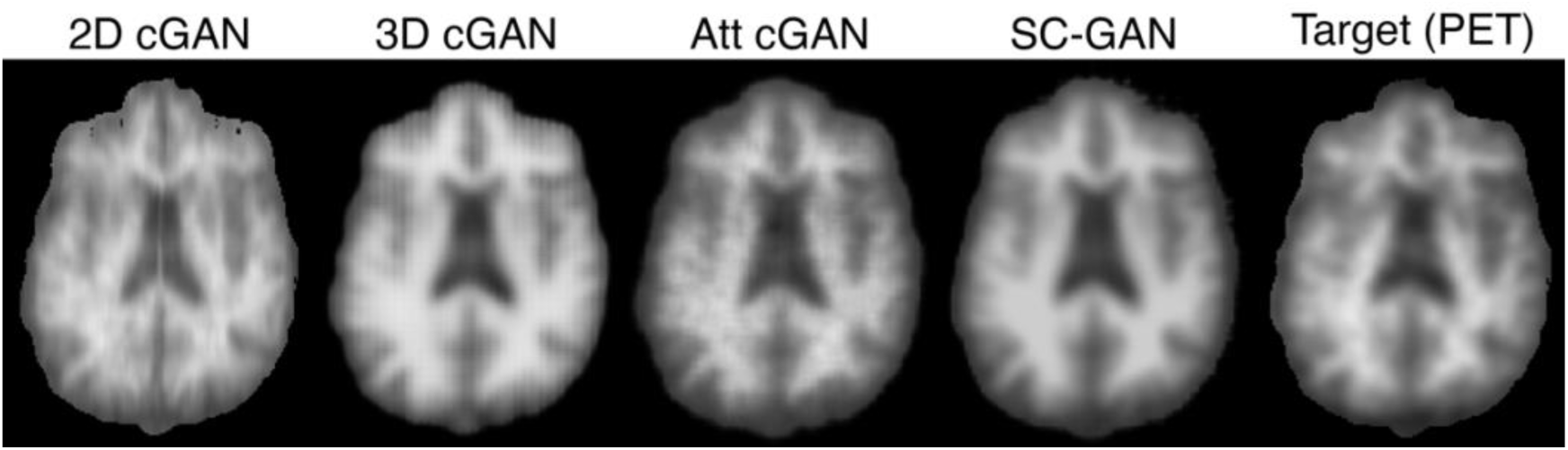
Qualitative assessment of PET synthesis task. Images are results of applying different GANs on T1w and FLAIR input images to predict Amyloid-beta PET. Target PET is also illustrated for comparison. Target image is normalized to [0 1] range for training and an equal color range of [0 1] are used for visualization.

**Figure 7.**
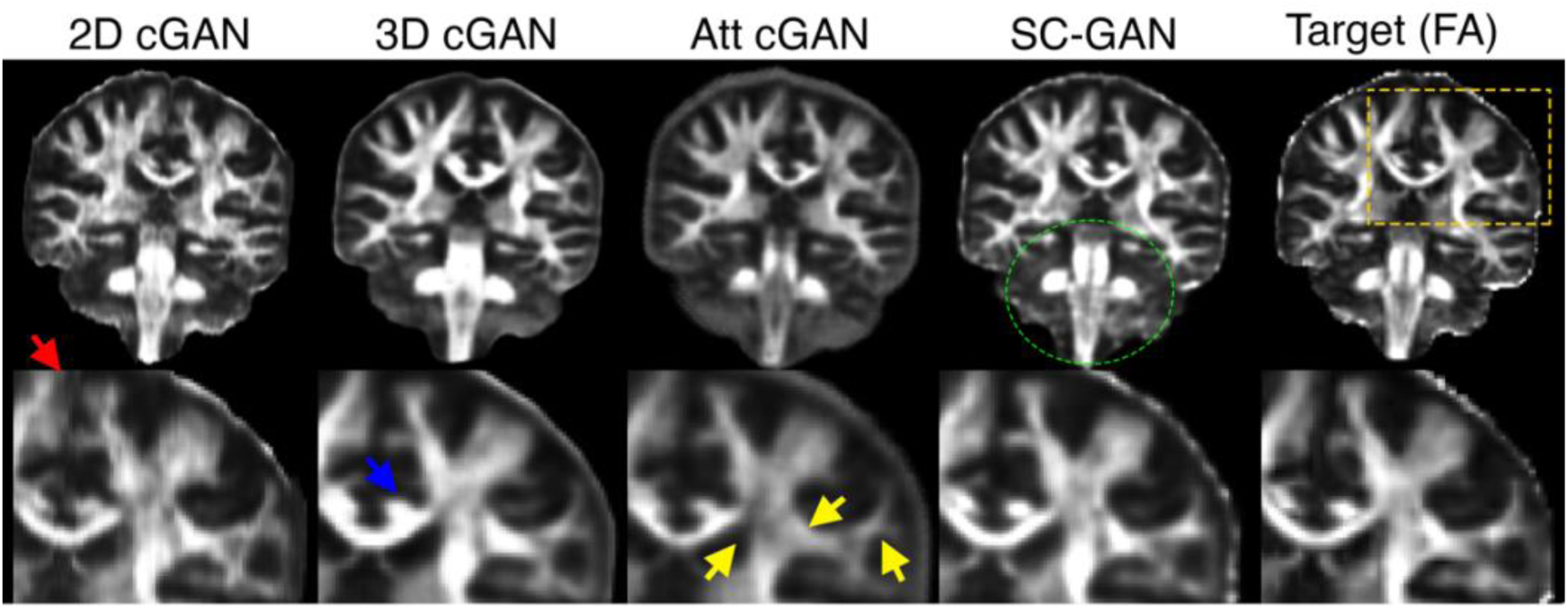
Qualitative assessment of fractional anisotropy (FA) synthesis task. Images are results of applying different GANs on T1w and FLAIR input images to predict FA. Target FA is also illustrated for comparison. An equal color range of [0 1] are used for visualization. Note that SC-GAN were able to synthesize FA in more detail in comparison with other networks. The 2D network demonstrated continuous distortion (red arrow), 3D cGAN resulted to an oversmoothed image (see blue arrow showing partial volume effect between fiber bundles of cingulum and corpus callosum). Attention cGAN failed to capture high intensity FA across the white matter (yellow arrows). Green dotted circle shows that, unlike other networks, SC-GAN was able to capture brainstem details.

**Figure 8.**
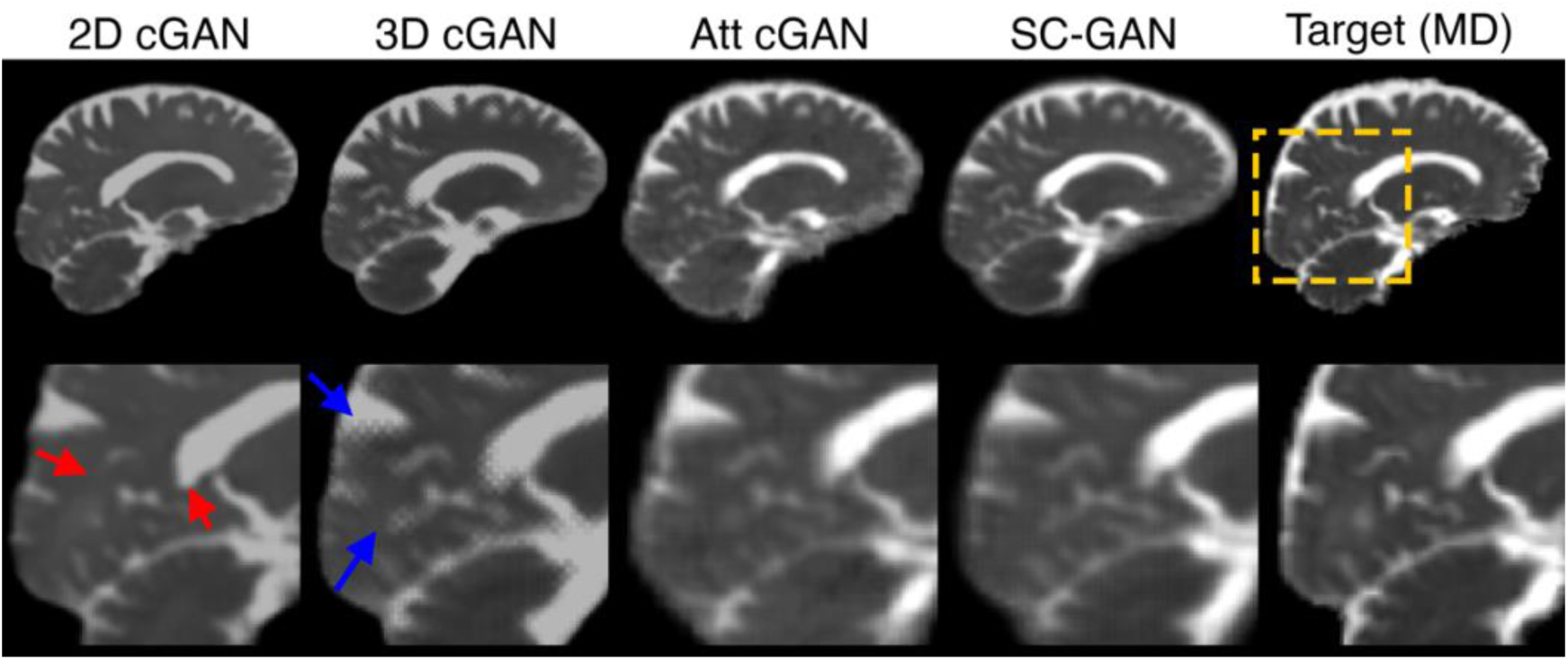
Qualitative assessment of mean diffusivity (MD) synthesis task. Images are results of applying different GANs on T1w and FLAIR input images to predict MD. Target MD is also illustrated for comparison. Target image is normalized to [0 1] range for training and an equal color range of [0 1] are used for visualization. Note that SC-GAN were able to synthesize MD in more detail in comparison with other networks. The 2D generated artificial sharp boundaries (red arrow) and 3D cGAN resulted to a large amount of striping artifact (blue arrow).

### Quantitative assessment

The learning curves show that all networks were successfully optimized, reaching the plateau within the range of the study epochs (**Figure 4**). 3D cGAN and SC-GAN networks showed a smooth and stable pattern in their optimization curve, while 2D cGAN and att GAN demonstrated a degree of fluctuation during the learning. The pattern of the learning curves across tasks was similar in SSIM and NRMSE. However, the PSNR was slightly different across tasks, with PET tasks resulted to the highest PSNR (**Figure 4**).

Regardless of the evaluation metric or the synthesis task, SC-GAN outperformed other networks, resulting to the lowest NRMSE and the highest PSNR and SSIM (**Figure 5** and **Table 1**). The NRMSE results showed that error of SC-GAN was 18%, 24% and 29% lower compared to 2D cGAN across FA, PET and MD tasks, respectively. Across all tasks, the 2D network resulted to the lowest performance.

**Table 1.**
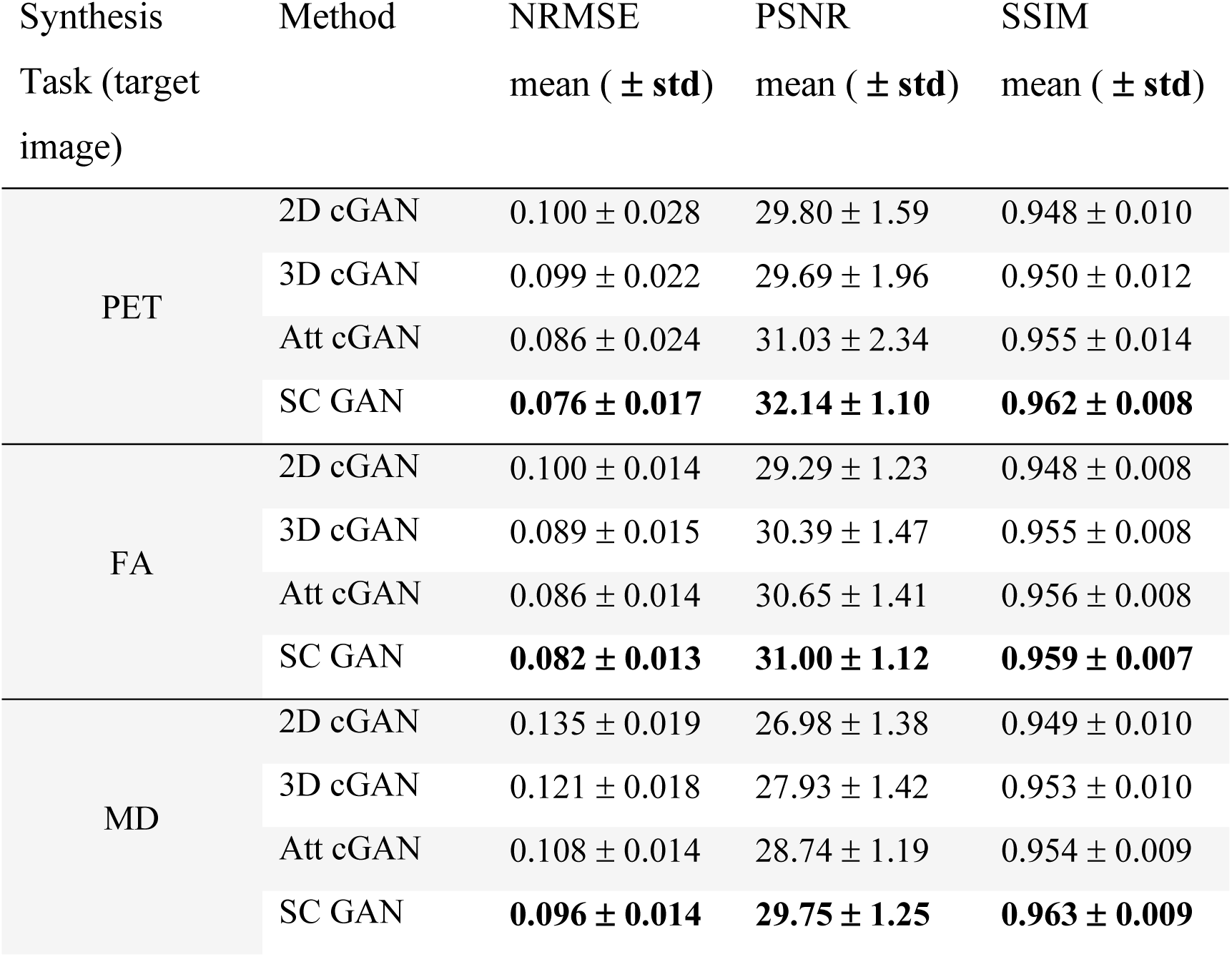
Comparison among different networks. Table shows statistic values of NRMSE, PSNR SSIM among test images after the networks reached the plateau and the hyperparameters were optimized. Statistically significant results are highlighted in bold font.

All 3D networks outperformed the 2D network, highlighting the importance of incorporating 3D information into deep learning networks. SC-GAN outperformed 3D cGAN and att GAN in all three tasks across all evaluation metrics. The increased performance of the SC-GAN was more evident in the PET task, followed by FA and MD tasks.

The ablation test showed that the major contributors to SC-GAN performance are the adversarial learning and the self-attention module, followed by B-rmse and spectral normalization modules (**Figure 9** and **Table 2**). Spectral normalization contributed to the stabilization of the SC-GAN training and feature matching loss contributed to generate synthesis result with more natural statistics at multiples scales.

**Table 2.** Ablation study of SC-GAN. Table shows ablation study of different components of SC-GAN on the Aβ PET synthesis task.

| Ablation study | NRMSE<br>mean ( ± std) | PSNR<br>mean ( ± std) | SSIM<br>mean ( ± std) |
| --- | --- | --- | --- |
| No self-attention | 0.118 ± 0.016 | 28.34 ± 1.200 | 0.939 ± 0.011 |
| No adversarial learning | 0.102 ± 0.018 | 29.72 ± 1.583 | 0.947 ± 0.012 |
| No brain area rmse loss | 0.092 ± 0.017 | 30.27 ± 1.627 | 0.953 ± 0.010 |
| No spectral normalization | 0.080 ± 0.017 | 31.57 ± 1.203 | 0.957 ± 0.010 |
| No feature matching | 0.078 ± 0.019 | 32.03 ± 1.174 | 0.960 ± 0.013 |
| SC-GAN | **0.076 ± 0.017** | **32.14 ± 1.100** | **0.962 ± 0.008** |

**Figure 9.**
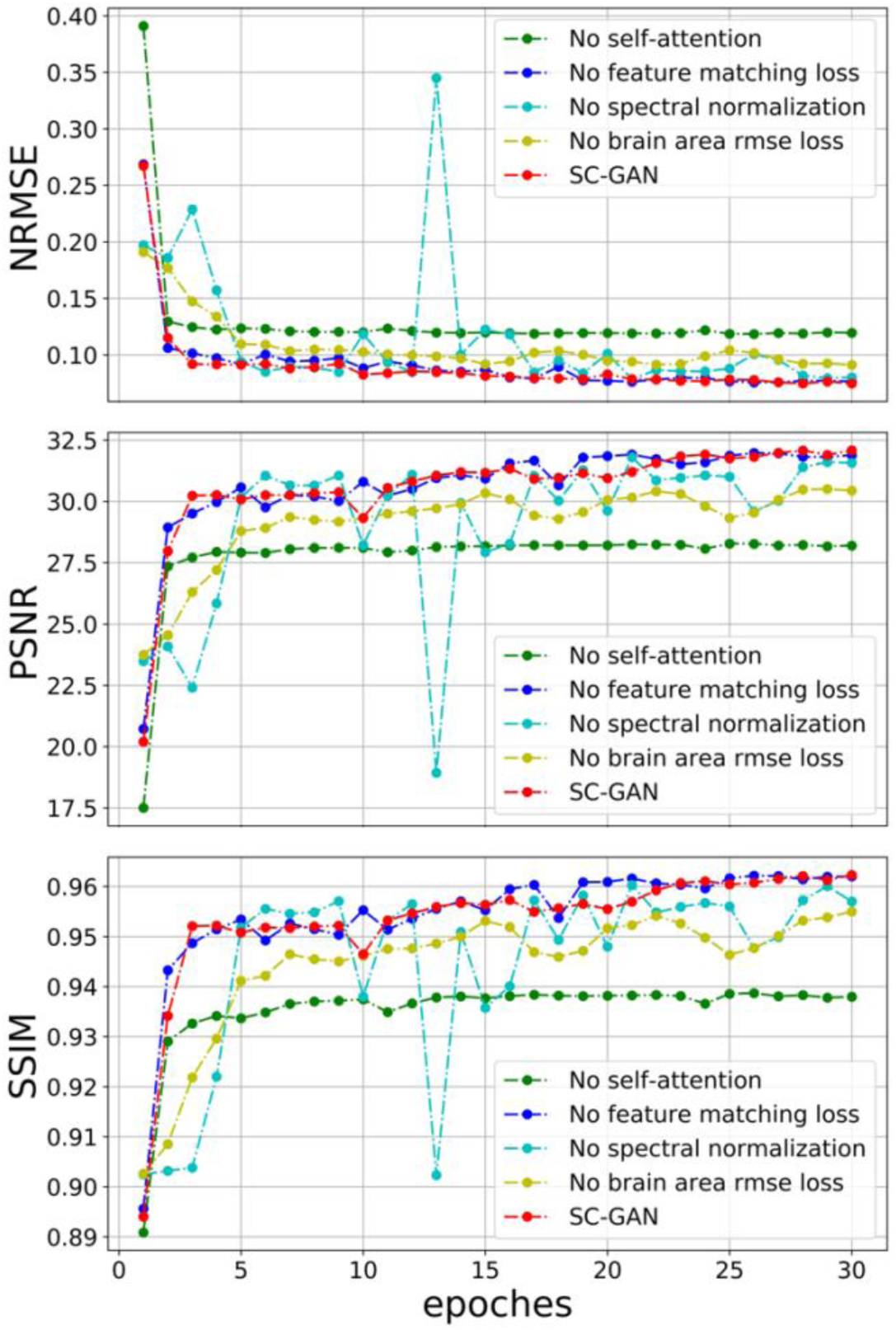
Ablation test across modules of SC-GAN. The SC-GAN with and without different network modules were assessed on the Aβ PET synthesis task and learning curves across different evaluation criteria are presented here. Plots demonstrate normalized mean square error (NRMSE), peak signal-to-noise ratio (PSNR) and structural similarity (SSIM). The self-attention module appeared to have the highest contribution to the achieved improvement, followed by spectral normalization and non-brain loss function exclusion.

### Qualitative assessment

Figure 6-8 compare the studied networks qualitatively. To assess the quality of the synthesis images in 3D, images were view across different plans: Axial images for PET synthesis (**Figure 6**), coronal images for FA synthesis (**Figure 7**) and sagittal images for MD (**Figure 8**). Because 2D cGAN was trained on the sagittal images, the sagittal view of the synthesized result provided the best result for the 2D network (e.g. MD task: **Figure 8**), while the axial and coronal views presented visual discontinuity and distortion (e.g. PET and FA tasks: **Figure 6** and **7**). Even at sagittal view, 2D GAN generated sharp artificial boundaries (ventricle boundaries in **Figure 8**). 3D network did not suffer from either of these shortcomings, presenting stable results across image dimensions.

SC-GAN results were visually closest to the ground truth data in comparison with other networks. In particular, SC-GAN was able to capture certain details in the image that were hidden to other networks. For example, structural boundaries at brain stem of the FA images were captured by SC-GAN (green dotted circle in **Figure 7**), but these details were smoothed out when other networks were used. Cingulum bundle (blue arrows, **Figure 7**) and superficial white matter (red arrow, **Figure 7**) were not generated with 3D cGAN and 2D cGAN, respectively. These details were successfully generated by SC-GAN. We also noted that att GAN failed to capture high intensity FA across the white matter (yellow arrows, **Figure 7**), whereas SC-GAN demonstrated a similar intensity profile as the ground truth. It should be noted that the SC-GAN also did not generated an exact match of the ground truth – artificial and incorrect features were observed. Results from MD synthesis (**Figure 8**) also showed that SC-GAN resulted to the generation of a map that is closest to the ground truth in comparison with other networks and contains higher degree of detail and less amount of artifact.

## Discussion

Here we presented an efficient end-to-end framework for multi-modal 3D medical image synthesis (SC-GAN) and validated it in PET, FA and MD synthesis applications. In order to design and optimize the network, we added a 3D self-attention module to the conditional GAN, which models the similarity between adjacent and widely separated voxels of the 3D image. We also employed spectral normalization and feature matching to stabilize the training process and ensure SC-GAN generate more realistic details (less artifacts). SC-GAN technique also allows multi-modal (multi-channel) 3D image input. We showed that SC-GAN significantly outperformed the state-of-the-art techniques, enabling reliable and robust deep learning-based medical image synthesis. SC-GAN is made available via https://github.com/Haoyulance/SC-GAN.

To obtain a generalized multi-modal 3D neuroimaging synthesis technique, SC-GAN incorporates adversarial learning, self-attention mechanism and stable learning strategy. SC-GAN network structure is demonstrated in **Figure 2**. The generator and discriminator are equipped with 3D self-attention modules, which can capture both short-and-long range dependencies for each feature vector during the learning process.

The self-attention feature makes SC-GAN a context-aware network, providing additional degree of freedom to the synthesis process. The ablation experiment conducted in this study showed that self-attention module contributed the most to the improvement of the conventional 3D GAN. Previous studies have shown that self-attention can be effective in other medical image analysis applications as well. Zhao *et al* (Zhao et al., 2020) combined object recognition network and self-attention guided GAN into one training process to handle tumor detection task. Li *et al* (Li et al., 2020) incorporated self-attention and auto encoder perceptual loss into convolutional neural network to denoise low dose CT.

While adding attention module improved the 3D cGAN, it provided less accurate results in comparison with SC-GAN that uses self-attention module. Att cGAN employs attention gate that filters the features propagated through the skip connections to enhance the feature maps at the upsampling phase. Since training process of Att cGAN is also guided by the attention gate module, network performance was better than 3D cGAN. Qualitative results also showed that Att cGAN can generated better results compared to 3D cGAN.

3D Medical image processing tasks often face dimensionality challenges, and GAN is no exception (Lundervold and Lundervold, 2019). 3D cGAN resulted to oversmoothed images in FA synthesis task and generated a large amount of striping artifact that resulted to blurring of the edges at PET and MD synthesis tasks. SC-GAN utilizes a series of regularization and stabilization techniques, namely feature matching loss, spectral normalization loss, L1 loss and brain area RMSE loss, allowing stable training on high dimensional input data (e.g. the input image size of N x 256 x 256 x 256 x 2 was used in this study).

The SC-GAN without adversarial learning resulted to a lower synthesis accuracy compared to the main implementation. SC-GAN without adversarial learning abandons the discriminator during the training phase. Since generator is a U-net like encoder-decoder (Çiçek et al., 2016; Ronneberger et al., 2015), the SC-GAN without adversarial learning is technically a 3D U-net with 3D self-attention module. The synthesis results of SC-GAN with and without the adversarial learning showed that the adversarial learning empowers the training process and could extend the plateau of the learning curve.

We incorporated a feature matching loss as part of the generator loss to stabilizes the training by forcing the generator to produce natural statistics at multiples scales. The discriminator takes target and synthesis images as inputs sequentially (**Figure 2**). Then, the cross-entropy loss is calculated to update the weights using a back-propagation approach. The feature matching loss uses the feature maps that are generated in the discriminator phase to produce similar output to the target image by minimizing the error associated with image spatial features.

The spectral normalization was used to stabilize the training process and prevent training from collapsing. Spectral normalization utilizes the Lipschitz continuity concept to impose constraint on the solution space (Hager, 1979) which stabilized SC-GAN training process (Miyato et al., 2018). Spectral normalization uses the Cauchy-Schwarz inequality on the Lipschitz continuity to bound the solution space, which stabilize the optimization.

Several recent works have used adversarial learning strategy for medical image synthesis (Li et al., 2020; Liu, 2019; Lundervold and Lundervold, 2019; Ouyang et al., 2019; Wang et al., 2019; Zhao et al., 2020). Most of the medical image synthesis and reconstruction works have been implemented using 2D or 2.5D input images (Li et al., 2020; Ouyang et al., 2019; Zhao et al., 2020). One drawback of 2D GAN is that the network can only utilizes one 2D image of axial, coronal or sagittal each time, and therefore, the synthesis 3D images present visual discontinuity, which appears similar to stripping artifact (**Figure 7** and **8**). To evaluate the benefits of 3D implementation, we compared the performances of 2D cGAN and 3D networks. We observed intensity discontinuity and distortion in the synthesis results of the 2D cGAN, which highlights the importance of utilizing 3D neural network implementation for medical image applications.

Recent works have shown that 3D GAN can be utilized to improve the accuracy of the medical imaging synthesis (Liu, 2019; Wang et al., 2019). To the best of our knowledge Wang et al first expanded the medical image synthesis GAN from 2D to 3D by utilizing 3D convolution and transposed convolution to achieve high-quality PET image estimation from low dose PET images (Wang et al., 2019). In order to rigorously assess SC-GAN, two existing 3D synthesis methods (3D cGAN and Att cGAN) were compared with SC-GAN. SC-GAN resulted to the highest performance and most stable learning curves (**Figures 4-5**).

It should be noted that while neuroimaging synthesis has drastically improved over the past five years, we do not think that synthesis can entirely substitute a given modality that is different in nature (for example PET). To assess the performance of image synthesis for detecting pathology in a cross-modal application, we estimated regional Amyloid update from synthesis PET and compared it with the ground truth PET (**Figure 10**). We noted a significant correlation between PET and synthesis PET across subcortical and cortical regions (*p*<0.0001 across all ten tested participants). Results were consistent across all test subjects, with correlation coefficient ranging from *r*=0.67 to *r*=0.95 (all with *p*<0.0001). While synthesis PET SUVR values were significantly correlated with those from ground truth PET, we observed that the error rate is higher when SUVR of the PET images are higher. These SUVR range corresponds to regions with high clinical value, reflecting neurodegenerative pathology (high Aβ uptake). Therefore, our results suggest that synthesis PET cannot substitute PET imaging, because pathological and clinically relevant molecular information in PET may not be detected by synthesizing PET that is obtained from MRI (which are mainly contain structural information). Nevertheless, this limitation does not damper the significance of medical image synthesis but calls for a careful design/application when image synthesis is used. For example, studies have shown that incorporating low-dose PET as synthesis input, reliable transformation can be achieved (Chen et al., 2019; Ouyang et al., 2019; Wang et al., 2019).

**Figure 10.**
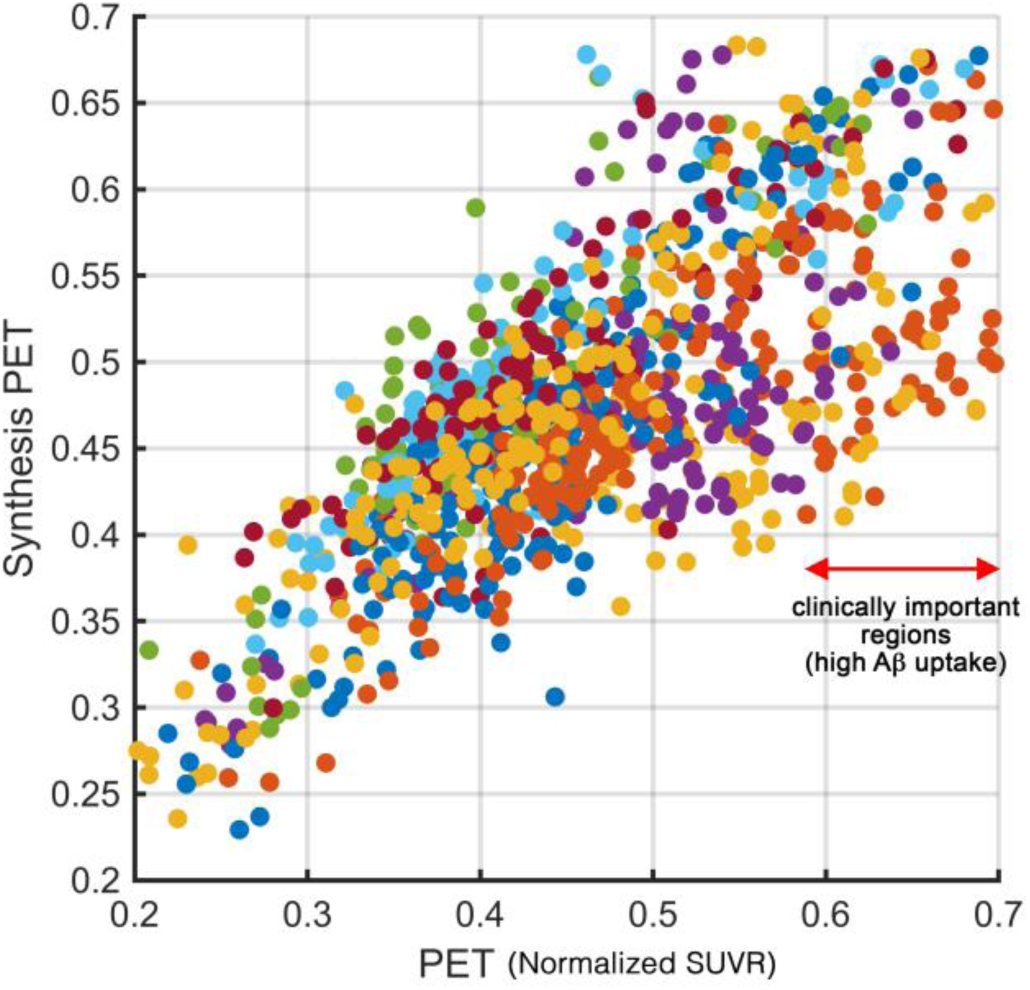
Correlation between PET and synthesis PET. Plot shows the correlation between Aβ standard uptake value ratio (SUVR) across subcortical and cortical regions of ten test participants (each color represents regions of each participants). PET images that were used for training were normalized using min-max normalization approach. Therefore, test PET images were also normalized using the same approach before comparison. Note that on region with high load of Aβ (shown with red arrow), the synthesis error is higher, suggesting that synthesis PET could not substitute PET imaging.

The focus of this work was on enabling multi-modal 3D neuroimage synthetization with GAN. The proposed method for this multi-modal 3D neuroimage synthesis (SC-GAN) was evaluated on the challenging task of PET and DTI synthesis to aid rigorous optimization of the network. SC-GAN is not intended to substitute PET with MRI-based PET synthesis. SC-GAN was designed and assessed to enable robust and stable multi-modal 3D neuroimaging synthesis. Future work will explore SC-GAN application. For example, SC-GAN may be used to combine MRI with low-dose PET to improve the efficacy of the existing techniques (Chen et al., 2019; Ouyang et al., 2019). We also expect that neuroimaging techniques with high number of repetitions such as functional and diffusion MRI (Ning et al., 2018) may benefit from SC-GAN, which is a future direction of our work.

## Acknowledgement

This work was supported by NIH grants: 2P41EB015922-21, 1P01AG052350-01, U54EB020406, USC ADRC 5P50AG005142. The content is solely the responsibility of the authors and does not necessarily represent the official views of the NIH.

## ADNI

Data collection and sharing for this project was funded by the Alzheimer’s Disease Neuroimaging Initiative (ADNI) (National Institutes of Health Grant U01 AG024904) and DOD ADNI (Department of Defense award number W81XWH-12-2-0012). ADNI is funded by the National Institute on Aging, the National Institute of Biomedical Imaging and Bioengineering, and through generous contributions from the following: AbbVie, Alzheimer’s Association; Alzheimer’s Drug Discovery Foundation; Araclon Biotech; BioClinica, Inc.; Biogen; Bristol-Myers Squibb Company; CereSpir, Inc.; Cogstate; Eisai Inc.; Elan Pharmaceuticals, Inc.; Eli Lilly and Company; EuroImmun; F. Hoffmann-La Roche Ltd and its affiliated company Genentech, Inc.; Fujirebio; GE Healthcare; IXICO Ltd.; Janssen Alzheimer Immunotherapy Research & Development, LLC.; Johnson & Johnson Pharmaceutical Research & Development LLC.; Lumosity; Lundbeck; Merck & Co., Inc.; Meso Scale Diagnostics, LLC.; NeuroRx Research; Neurotrack Technologies; Novartis Pharmaceuticals Corporation; Pfizer Inc.; Piramal Imaging; Servier; Takeda Pharmaceutical Company; and Transition Therapeutics. The Canadian Institutes of Health Research is providing funds to support ADNI clinical sites in Canada. Private sector contributions are facilitated by the Foundation for the National Institutes of Health (www.fnih.org). The grantee organization is the Northern California Institute for Research and Education, and the study is coordinated by the Alzheimer’s Therapeutic Research Institute at the University of Southern California. ADNI data are disseminated by the Laboratory for Neuro Imaging at the University of Southern California.

